# Cortical time-course of evidence accumulation during semantic processing

**DOI:** 10.1101/2022.06.24.497472

**Authors:** Gayane Ghazaryan, Marijn van Vliet, Lotta Lammi, Tiina Lindh-Knuutila, Sasa Kivisaari, Annika Hultén, Riitta Salmelin

## Abstract

Our understanding of the surrounding world and communication with other people are tied to mental representations of concepts. In order for the brain to recognize an object, it must determine which concept to access based on information available from sensory inputs. In this study, we combine magnetoencephalography and machine learning to investigate how concepts are represented and accessed in the brain over time. Using brain responses from a silent picture naming task, we track the dynamics of semantic processing, and show that the brain gradually accumulates semantic information before eventually reaching a plateau. The timing of this plateau point varies across individuals, indicating differences in the temporal domain of semantic processing, a new dimension of individual variability in the faculty of language.

## 1. Introduction

Concepts are fundamental building blocks of our understanding of the world and communication with others. Brain regions associated with semantic knowledge have been extensively studied (Binder & Desai, 2011; Binder, Desai, Graves, & Conant, 2009; Lambon Ralph, Jefferies, Patterson, & Rogers, 2017; Liuzzi, Aglinskas, & Fairhall, 2020), yet the mechanisms of how concepts are accessed in the brain are still not well understood. Object recognition is a common task that requires accessing concepts, and it is a prime target for experimental research on semantic processing. Despite how rapidly we can recognize an object, the process likely consists of multiple phases (Levelt, Praamstra, Meyer, Helenius, & Salmelin, 1998), involving the interplay between visual and semantic properties (Clarke, 2019) and the emergence and accumulation of information over time (Clarke, Devereux, Randall, & Tyler, 2015; Contini, Wardle, & Carlson, 2017). In this work, we track and examine the dynamic progression of semantic processing in the human brain over the course of visual object recognition. We hypothesize that the brain accumulates semantic information before accessing the appropriate semantic concept and identifying the object.

Object recognition is essential in everyday functioning and, indeed, it has received great interest in human neuroscience (see Clarke, 2019; Mahon & Caramazza, 2009; Martin, 2007; Wardle & Baker, 2020, for reviews). The underlying process is thought to progress from a focus on low-level visual features to a focus on complex semantic representations (Clarke, 2019). Semantic knowledge models (Bruffaerts et al., 2019; Harris, 1954; Joos, 1950; Osgood, Suci, & Tannenbaum, 1978) together with machine learning methods (Mitchell et al., 2008; Palatucci, Pomerleau, Hinton, & Mitchell, 2009) have been utilized to link brain activation patterns and semantic processing. Applying such methods to time-sensitive neuroimaging data, semantic processing has been shown to follow a coarse-to-fine progression (Clarke, 2019). Coarse semantic categories, but not individual concepts, can be discriminated based on earlier brain response patterns; by around 150 ms, it is possible to decode categories of objects (Clarke, 2019). Individual concepts, however, can be decoded only at later time points, around 300–450 ms after stimulus onset (Chen et al., 2016; Clarke et al., 2015; Rupp et al., 2017; Sudre et al., 2012).

Previous studies, which have focused on decoding at isolated time windows in sequence, have revealed a pattern of increasing decoding accuracy up to a peak, followed by a gradual decrease in accuracy (Clarke et al., 2015; Rupp et al., 2017; Sudre et al., 2012). Often, such studies report the earliest time-point when decoding is significantly above chance. However, there are multiple factors that influence the earliest time window when decoding is considered successful, so it can be a noisy indicator. Instead, we consider stabilization of the decoding accuracy to be more informative of the progression of semantic processing.

Furthermore, cross-temporal decoding (decoding information in one time window with models trained on other windows) has shown that the underlying brain activation patterns evolve rapidly following stimulus onset, with some generalization of similar encoded information across nearby time windows (e.g., Carlson, Tovar, Alink, & Kriegeskorte, 2013; Grootswagers, Wardle, & Carlson, 2017). Such decoding behavior suggests an interplay between accumulation and maintenance of information (Contini et al., 2017; Ploran et al., 2007). Indeed, for brain-level object recognition we need to take into account all information gathered up to a certain time point, instead of limiting the view to a sequence of single snapshots, analyzed in isolation.

To investigate how the brain processes information before reaching a decision, we use MEG (magnetoencephalography) brain response data from a picture viewing experiment, in which participants were shown pictures of objects and asked to silently identify them (Figure 1A). This task focuses on object perception through to concept access (Levelt et al., 1998), and excludes later processes involving phonological forms and speech production. To ensure invariance of low-level visual features and emphasis on semantic properties, we employ averaged brain responses to multiple different exemplars of the same concept. We contrast two approaches for decoding semantic representations: the traditionally used sliding window of fixed-length and a novel cumulative modeling approach (Figure 1B) that widens the window at each time step. We demonstrate that, indeed, the brain gradually accumulates semantic information with eventual stabilization of decoding accuracy and access to object identity.

**Figure 1:**
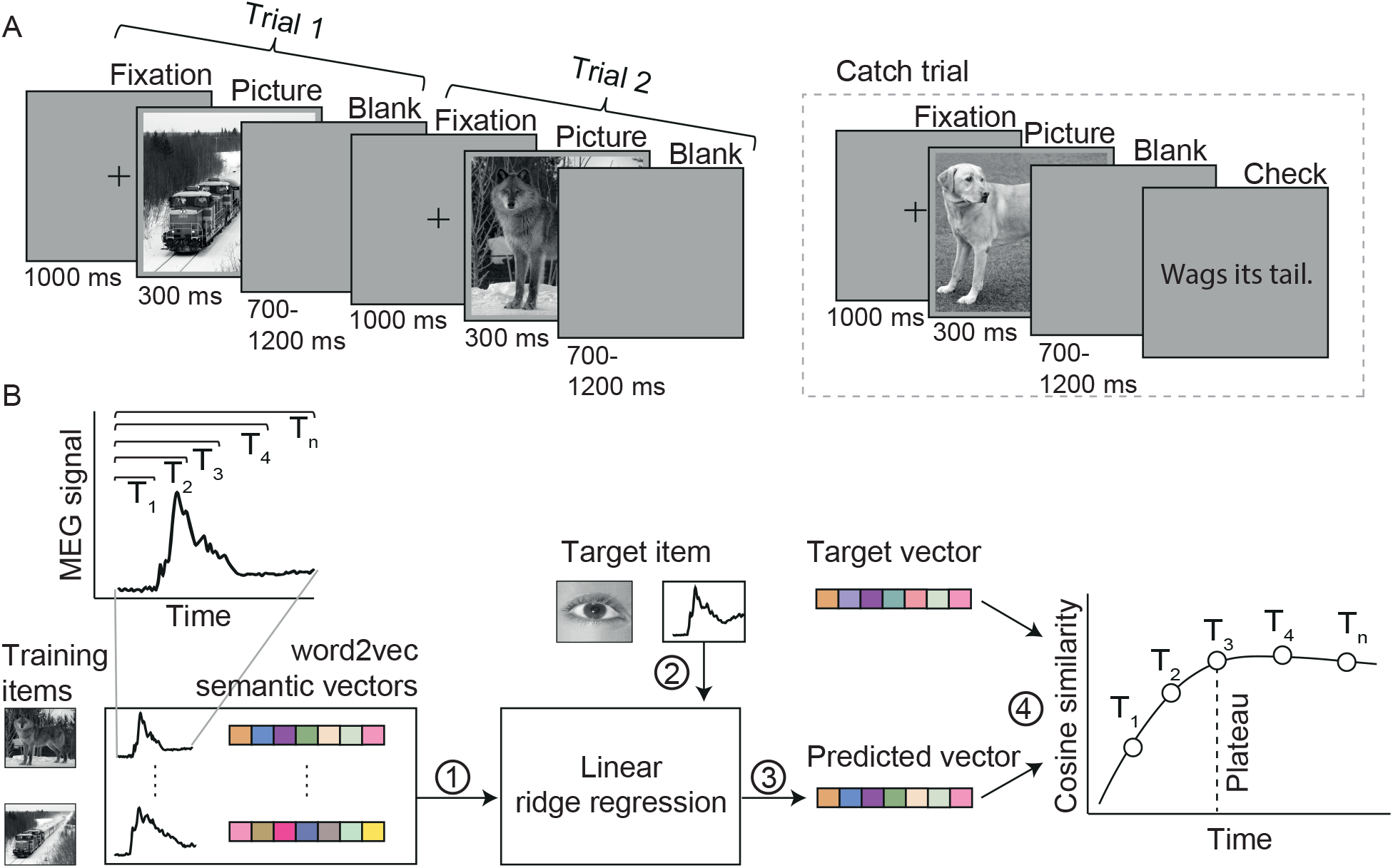
**A**. Experimental paradigm. In each trial, participants were presented with a picture of an object and asked to identify it; randomly occurring catch trials ensured compliance. **B**. Overview of the brain-to-semantics mapping method. Linear regression models were (1) trained on a set of brain response-semantic vector pairs and (2) tested on the brain response to one left-out concept, (3) yielding a predicted vector. This was (4) compared to the semantic vector of the target concept using cosine similarity. This procedure was repeated for all concepts. The same procedure was performed for each instance of the cumulative window. The dashed line corresponds to the point after which prediction-target similarity no longer increases (plateau point). Semantic vectors were created using the word2vec algorithm on a Finnish text corpus.

## 2. Results

### 2.1. Validation of MEG data and semantic model

We trained a regression model to predict the semantic representation of a concept from the average MEG response to a set of different exemplars of that concept. The word2vec algorithm was used to represent each concept as a vector in semantic space (Mikolov, Sutskever, Chen, Corrado, & Dean, 2013), based on co-occurrence statistics in a Finnish text corpus. We used a linear-regression based zero-shot decoding approach, where the model is trained on a set of brain response-semantic vector pairs and tested on brain responses to concepts it had not been trained with (Palatucci et al., 2009). Predictions were then evaluated by matching them against a list of possible concepts (the target and 99 other concepts not used as stimuli). The decoding was evaluated by leave-one-out rank accuracy (Markowitz, Schmidt, Burlina, & Wang, 2017).

Similar approaches have been successfully used in earlier studies of semantics in the brain (Hultén et al., 2021; Kivisaari et al., 2019; Mitchell et al., 2008; Sudre et al., 2012). We first verified the suitability of the decoding approach for each participant by checking that the model could successfully decode the concepts from the entire brain response pattern at 0–1000 ms after stimulus presentation. The modelling approach was shown to be valid for all participants (Figure 2A), with mean rank accuracy ranging between 74% and 89% (to significantly exceed chance level, the required rank accuracy was determined to be greater than 68%, based on a permutation test).

**Figure 2:**
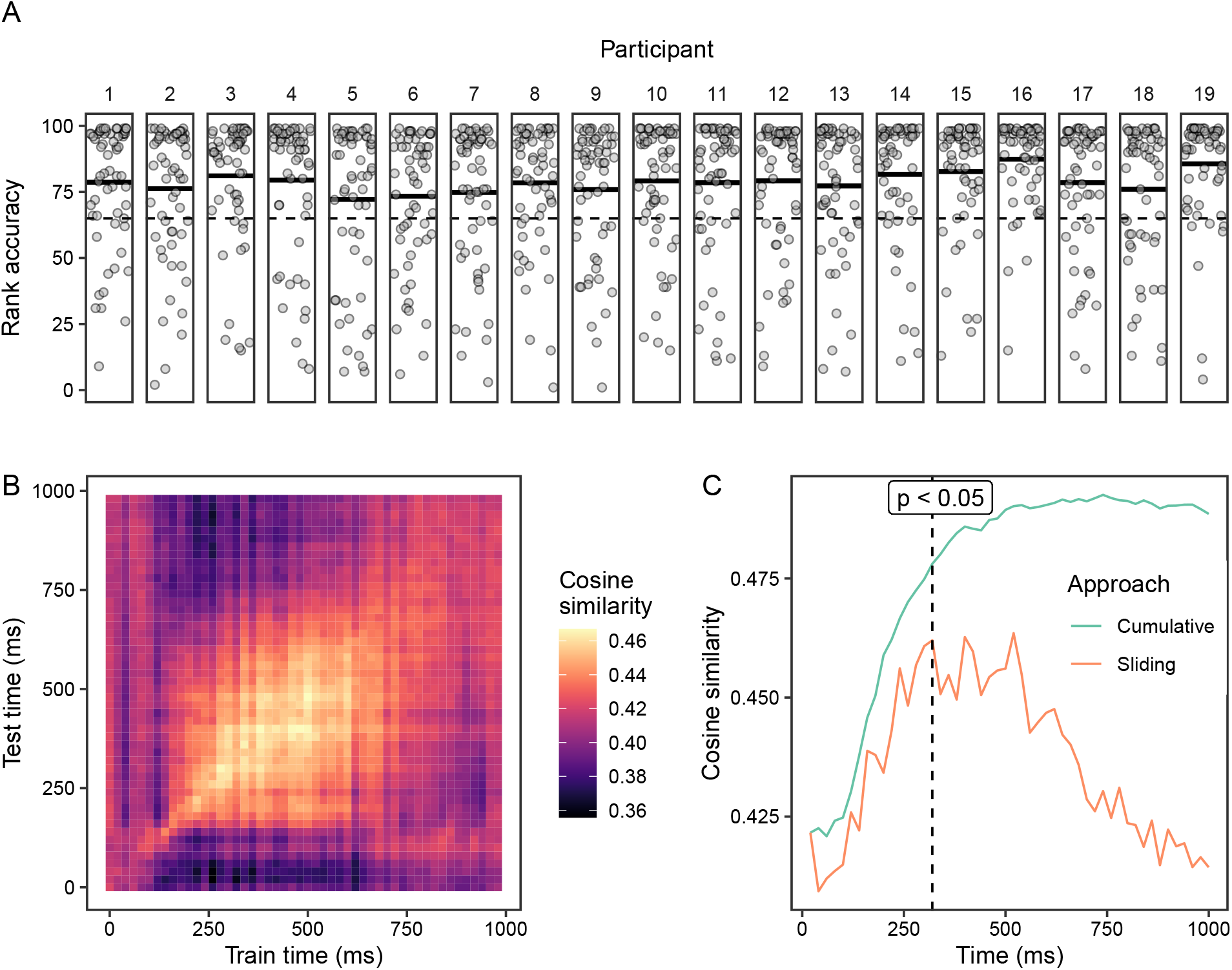
**A**. Zero-shot decoding accuracy based on leave-one-out cross-validation for each participant. The model predictions were evaluated in relation to 99 other concepts not included as stimuli, using the rank of the predicted concept. Dots indicate target concepts, solid black lines indicate participant-level averages, and the dashed line corresponds to chance level performance (as determined through a permutation test). **B**. Temporal cross-decoding results on grand average data. Here, models were trained on one 20 ms time window and tested on another 20 ms time window. The color corresponds to the prediction-target cosine similarity averaged over all targets. **C**. Similarity between predictions and targets over time (cosine similarity) on grand average data, using two different types of models: a sliding window of length 20 ms without overlap (i.e. 180 ms on the x-axis indicates the model was trained on 160–180 ms) and a cumulative window with width increasing with 20 ms increments (i.e. 180 ms on the x-axis represents models trained on 0-180 ms after stimulus onset). The dashed line corresponds to the point after which the prediction-target similarity difference between cumulative and sliding window approaches is significant (as determined through a permutation test).

### 2.2. Temporal progression of semantic processing

Next, we examined decoding across different time windows to uncover possible similarities of encoded semantic information between them. The models were trained on data from one time window and tested on another time window; this analysis was performed on MEG data averaged across participants. The resulting temporal cross-decoding patterns, averaged over all target concepts, revealed information generalization between 200 and 750 ms (Figure 2B).

We examined the underlying reason for this generalization by considering two alternatives: either the semantic progression finishes by the beginning of this period and the information is maintained, or accumulation of information continues throughout this period. We compared two different types of models with (1) sliding fixed-length and (2) cumulative windows. We expected the latter type of model to show a pattern of increasing prediction-target similarity, with an eventual plateau corresponding to the point after which there is no further improvement from including data from more time windows. We defined these plateau points by the time point after which there was less that 5% increase in prediction-target similarity. We interpreted this as the time after which little more semantic information could be decoded and object identification was reached.

Models with a fixed-length sliding window indicated that model predictions, averaged over all concepts, were closest to targets between 300 and 500 ms (Figure 2C). All time windows in this range resulted in similar performance. In contrast, the cumulative approach, which includes all preceding time points, reached its highest prediction-target similarity at 500–700 ms after stimulus presentation, after which the similarity no longer increased (Figure 2C). Furthermore, the cumulative window approach reached higher similarity values than the sliding window approach. This difference was statistically significant after 320 ms (based on a permutation test (*p <* .05). Analysis at the level of individual participants (Figure 3A) indicated that the mean plateau points of the cumulative models (after which the similarity value did not increase) were significantly later than the peaks of the sliding-window approach, *t*(18) = 6.997, *p <* .001.

**Figure 3:**
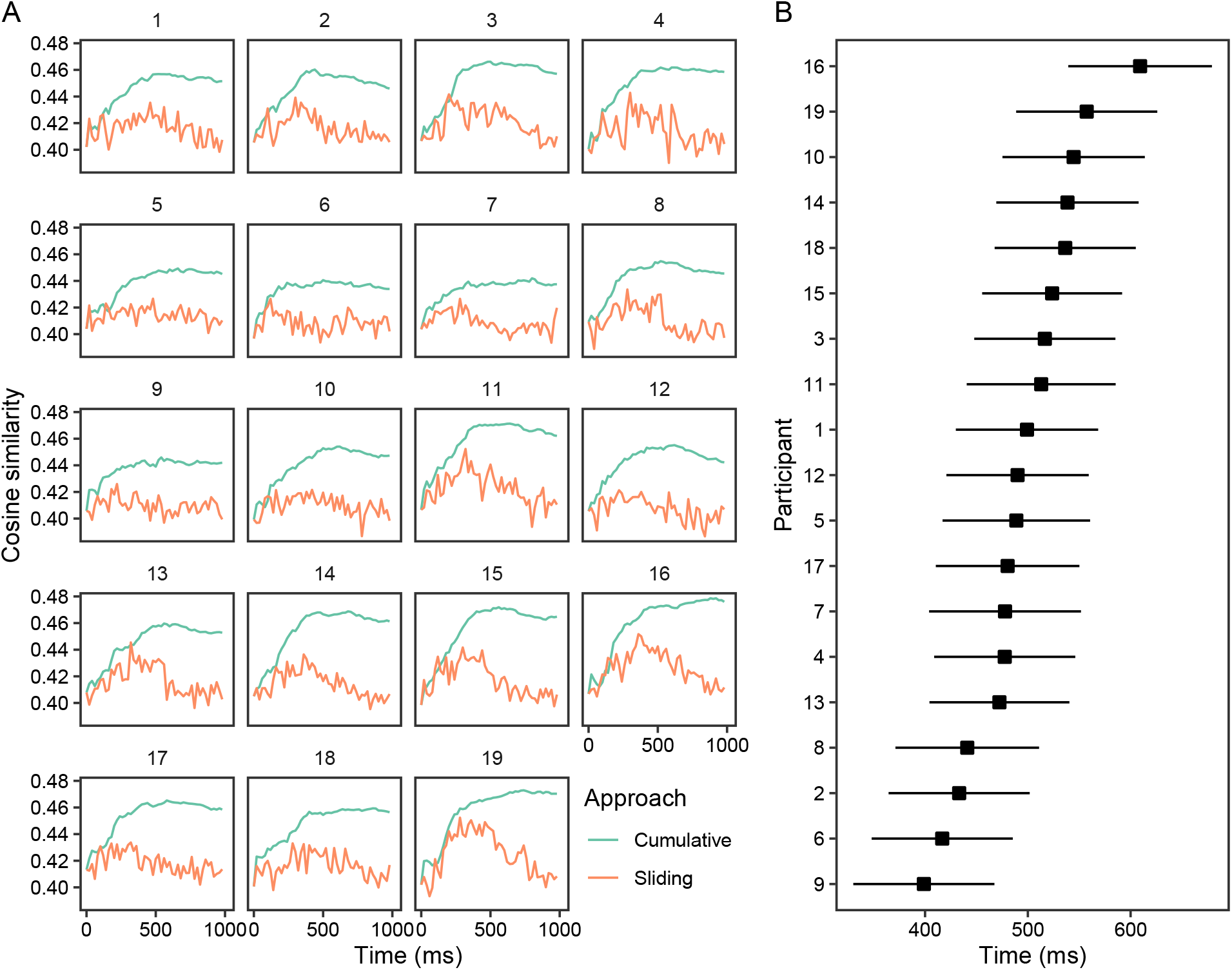
**A**. Similarity between predictions and targets over time in individual participants (cosine similarity). All participants displayed increasing prediction-target similarity as a function of time with an eventual plateau. Plateau points (for cumulative models) and peaks (for sliding fixed-length models) were defined as the time point after which prediction-target similarity no longer increased by more than 5%. **B**. Plateau points were modeled using a generalized linear mixed model, with fixed effects of participant and random intercepts for concepts. Depicted are the mixed model results for these plateau points showing their estimated marginal means and 95% confidence intervals.

### 2.3. Individual differences

As people have unique life experiences and different associations between concepts, the semantic processes underlying understanding of the world likely vary across individuals. We used the cumulative window model to investigate and compare the progression of semantic processing of each participant. All participants displayed increasing prediction–target similarity as a function of time with an eventual plateau (Figure 3A). We focused on the variation of the point of plateau between individuals. A generalized linear mixed model (with random intercepts for each concept) showed significant variation between participants’ plateau points (*F*(18, 981.90) = 2.29, *p* = .002) ranging from 400 to 600 ms (Figure 3B).

### 2.4. Representational similarity analysis

To investigate the brain areas involved in the information accumulation processes, we performed representational similarity analysis (RSA) between concept similarity in the brain (the brain-level concept-to-concept dissimilarity matrices at different time windows in sliding and cumulative windows; lower sections of Figure 4A and B) and concept similarity in the semantic model (the semantic model concept-to-concept dissimilarity matrix). We observed the highest RSA values (Spearman’s rank correlation coefficient) in the occipital regions of both hemispheres. Incrementally adding time windows did not change the regions where the RSA values were highest. The 0–400 ms to 0–700 ms windows showed the highest correlations, while correlations decreased slightly when considering the entire 0–1000 ms window. This timing aligned with the decoding results described above.

**Figure 4:**
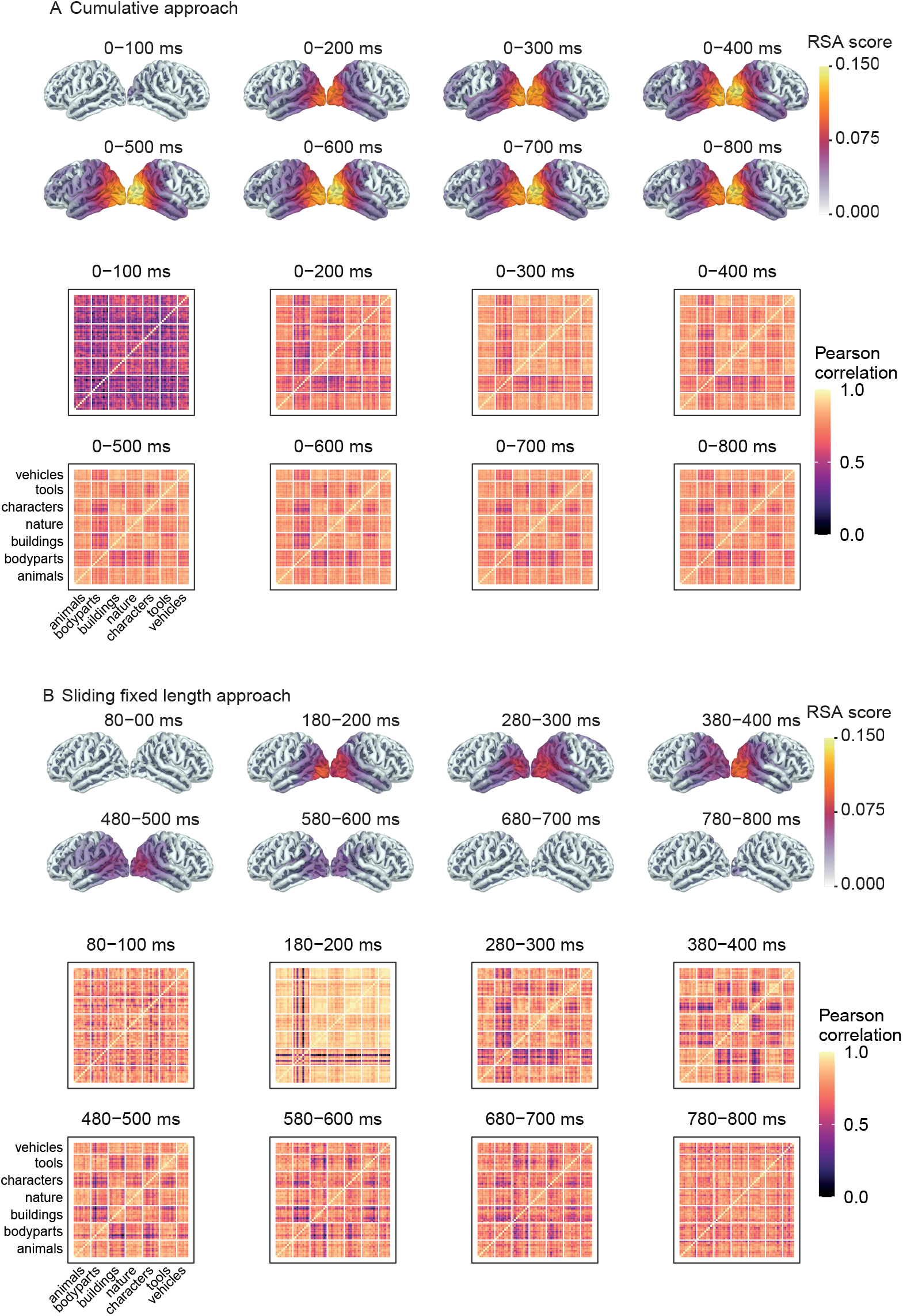
RSA maps illustrating the statistically significant clusters for different time windows, with the sliding fixed-length approach and (**B**) the cumulative approach. RSA values correspond to Spearman’s rank correlation coefficient between concept-to-concept brain dissimilarity matrices and the corpus-based model dissimilarity matrix. Significance was determined using a cluster permutation test. Below the RSA maps are concept-to-concept brain signal dissimilarity matrices (sensor-level) plotted over time, averaged across all participants. The RSA score corresponds to 1 *− r* where *r* is the pairwise Pearson correlation coefficient between brain responses to different concepts over time.

## 3. Discussion

Our high zero-shot decoding accuracy results using the entire 1000 ms of the brain response align with previous work, indicating that information related to semantic processing was successfully captured. We were further able to track the progression of semantic processing by decoding the neural correlates throughout this time window. The results from the sliding window approach indicated that there was information relevant to semantic processing between 250 and 600 ms after stimulus onset, which matches the timing reported in previous work (Clarke et al., 2015; Giari, Leonardelli, Tao, Machado, & Fairhall, 2020; Rupp et al., 2017; Simanova, van Gerven, Oostenveld, & Hagoort, 2010; Sudre et al., 2012). Temporal cross-decoding indicated that there was also generalization in the brain signal during this time. Contini et al. (2017) have suggested two possible reasons for such generalization: the encoded semantic representation is maintained throughout this period, or there are ongoing overlapping processes of differing duration, such that some information is maintained while the representation is enriched with further accumulation.

From around 320 ms onward, the cumulative approach resulted in predictions that are significantly closer to targets than the independently considered sliding windows. This indicates that at 320 ms, the process of object recognition had not yet finished, and information accumulation continues until the plateau at around 600 ms, rather than there being only maintenance of the object representation. These results imply that semantic processing of pictured objects occurs by information accumulation.

By using static images and multiple instances of each concept, we hoped to reduce the effects of mere stimulus characteristics on the observations and highlight neural effects related to processing of the concept. By demonstrating information accumulation, our work brings relevant new information to complement earlier studies on semantic access in general (Clarke et al., 2015; Deniz, Nunez-Elizalde, Huth, & Gallant, 2019; Huth et al., 2016; Huth, Nishimoto, Vu, & Gallant, 2012; Just, Cherkassky, Aryal, & Mitchell, 2010; Kivisaari et al., 2019; McCartney, Martinez-del Rincon, Devereux, & Murphy, 2019; Pereira et al., 2018; Shinkareva et al., 2008) and especially its temporal progression (Clarke et al., 2015; Hultén et al., 2021; Rupp et al., 2017; Sudre et al., 2012).

RSA indicated that occipital areas were relevant to semantic processing of pictures and showed patterns in accordance with the decoding approach. Interestingly, brain regions that are consistently reported in studies of picture naming (Ala-Salomäki, Kujala, Liljeström, & Salmelin, 2021), such as the left temporal and left parietal cortices, did not strongly account for semantic relationships between target concepts. However, our results align with Simanova et al. (2010), who suggested that the predominance of occipital areas may be due to inherent visual similarities between semantically similar objects. In other words, the appearance of an object is tied to its semantic meaning, so it is unsurprising that brain regions related to visual processing emerged in the RSA analysis. While it is possible that silent identification of the pictures did not activate the phonological form of the concept as strongly as an overt naming task would have done, the lack of involvement of the typical language areas may be a reflection of the fact that semantic similarity is not mirrored in phonological similarity (for example ‘cat’ and ‘dog’ are semantically near but phonologically distant).

We were further able to discern the end point of the progression of information accumulation, and we propose that it could be interpreted as the time point when object identification has been reached. On group-level data, the mean plateau time across concepts was around 500 ms. This, coupled with the generalization shown between 200 and 750 ms, indicates that there is an interplay between accumulation of information (preceding plateau) and maintenance (following plateau). The timing of the plateau point varied between participants, with the mean ranging from approximately 400 to 600 ms.

The fact that people agree on names of objects and can communicate about them indicates that there are commonalities in semantic understanding. However, as each person has a unique life experience, the underlying semantic processes are also likely to vary. Previous studies have reported inter-individual variation in behavioral measures of naming speed (Shao, Roelofs, & Meyer, 2012), neural correlates of semantic representation (Alfred, Hillis, & Kraemer, 2021), and gaze-behavior measures of visual salience (de Haas, Iakovidis, Schwarzkopf, & Gegenfurtner, 2019). Individual variation has also been indirectly investigated through cross-decoding between individuals. This is performed by training models on data from one or more individuals and testing on data from another individual (Just et al., 2010; Shinkareva, Malave, Mason, Mitchell, & Just, 2011; Shinkareva et al., 2008). Generally, such cross-decoding has been less accurate than within individual decoding. As these studies predominantly used imaging methods that favor high spatial precision over temporal precision, the results likely indicate individual variation in the cortical areas involved in language processing, the existence of which has been known since early studies (Ojemann, Ojemann, Lettich, & Berger, 1989). Individual differences in the temporal domain of semantic processing have also recently been indicated by Rupp et al. (2017), who reported individual variation in the windows when decoding performed best, suggesting differences in the progression of semantic understanding. Here, we have identified another dimension of variation in semantic processing: time until stabilization of semantic representations.

We have presented here a new perspective on the temporal dynamics of semantic understanding that opens future avenues of research. These include continued focus on individual variation, adapting to other modalities such as spoken or written words, and investigating concept processing in context by using more naturalistic stimuli such as sentences or stories. Such research will bring us towards a more complete model of language within the brain.

## 4. Materials and methods

### 4.1. Participants

20 native speakers of Finnish (females/males 10/10, age range 20-27, mean age 22) participated in the study. All participants were right handed (Edinburgh handedness questionnaire (Oldfield, 1971)) and had normal or corrected to normal vision. The study was approved by the Aalto University Research Ethics Committee and participants provided written informed consent prior to their participation. Data from 1 participant was excluded due to technical issues with the MEG recordings, leaving data from 19 participants in the final analysis.

### 4.2. Stimuli and procedure

Stimuli consisted of 300 grayscale photographic images of 60 concrete Finnish nouns (five different depictions of each). To minimize the effects of low-level visual features on the neural responses, there were five different images depicting each concept, the responses of which were then averaged. The nouns belonged to 7 different categories: animals, body parts, buildings, nature, human characters, tools/artifacts, and vehicles. Details of the nouns are presented in Appendix A. There were nine concepts in each category except for vehicles, which had six concepts. In the experiment, the concepts were also presented in written and auditory forms in separate trials; the responses from those trials were not analyzed in this study.

Participants were tasked with viewing each picture and silently identifying and thinking about the depicted object. The stimuli were presented at a size of 106 *×* 106 mm on a screen 140 cm from the participants’ eyes, corresponding to a visual angle of 4.3°. Each trial started with a fixation cross displayed for 1000 ms. The picture was then shown for 300 ms, follzowed by a blank screen for a randomized duration of 700–1200 ms (Figure 1A).

Measurements for each participant took place over 3 sessions on different days. Overall, each concept was presented 18 times. To ensure that participants remained engaged during the experiment, we included comprehension tasks after 10% of trials. In these tasks, participants used optical response pads to indicate whether or not a written description was characteristic of the previously shown concept. As these tasks occurred after the trials, they did not interfere with the responses, thus all trials were included in the analysis.

### 4.3. Data acquisition

MEG measurements were conducted at the Aalto NeuroImaging MEG Core (Aalto University, Espoo, Finland) using a Vectorview whole-head MEG system (MEGIN (Elekta Oy), Helsinki, Finland). The system has 306 sensors (204 planar gradiometers, 102 magnetometers). The head position was continuously tracked during the experiment by 5 head position indicator (HPI) coils placed at known locations with respect to identifiable anatomical landmarks. Eye movements and blinks were captured using 2 electrode pairs (one pair positioned above and below the left eye, the other in the corner of each eye). The recording was bandpass-filtered at 0.03–330 Hz and sampled at 1000 Hz. Anatomical MRIs were obtained using Siemens Magnetom Skyra 3.0 T MRI scanner with a T1-weighted MP-RAGE sequence at the Aalto NeuroImaging Advanced Magnetic Imaging (AMI) Centre.

### 4.4. Data preprocessing

MEG data was first visually inspected and noisy channels were identified. External sources of noise were then removed using spatiotemporal signal space separation (tSSS; Taulu & Simola, 2006) with Elekta Maxfilter software (MEGIN Oy, Finland). For each participant, data from different sessions was transformed to the same head position. All subsequent analysis was performed using the MNE-Python software package (Gramfort, 2013). The data was low-pass filtered at 40 Hz and split into 1200 ms epochs, the first 200 ms of which was the pre-stimulus baseline interval. To reduce contamination related to heartbeats, eye movements and blinks, we performed independent component analysis (ICA). In order to minimize the effect of slow drifts on ICA decomposition, we used continuous data high-pass filtered at 1 Hz (Jas et al., 2018). Components corresponding to heartbeats, eye movements and blinks were visually identified and excluded from epochs. Epochs corresponding to the same concept were then averaged, the time period between 0 and 1000 ms was extracted, and the signal was downsampled to create 20 ms bins. Only data from the gradiometers was used in the final analysis. This resulted in a matrix with 60 concepts *×* 204 channels *×* 50 time points for each participant.

We computed the source-level estimate of the average response to each concept using minimum norm estimates (MNE; Gramfort, 2013). Anatomical MRIs were used to reconstruct the cortical surface of each participant applying the FreeSurfer software package (Dale, Fischl, & Sereno, 1999; Fischl, 2012; Fischl, Sereno, & Dale, 1999). We used a single-layer boundary element model (BEM) with an icosahedron mesh of 2562 vertices in each hemisphere. When computing the inverse solution, a loose orientation constraint of 0.3 and depth weighting parameter of 0.8 were used. An empirical noise-covariance matrix was computed based on the pre-stimulus 200 ms interval to all concepts. To prepare the data for group-level analysis, participant-level source estimates of each concept were morphed to the FreeSurfer standard template brain (fsaverage).

### 4.5. Semantic space

Vector representations for the stimuli were obtained using the word2vec tool with skip-gram architecture and negative sampling algorithm (Mikolov et al., 2013). Each concept was represented as a vector of length 300 which defines its location in semantic space. The components of a vector are based on word co-occurrence statistics in large text corpora. Here co-occurrence was considered to take place when a word appeared within a window from 5 words before to 5 words after the word corresponding to the concept of interest. The corpus used was the Finnish Internet Parsebank (Luotolahti, Kanerva, Laippala, Pyysalo, & Ginter, 2015), which is based on a large sample (1.5 billion words) of Finnish language websites.

### 4.6. Machine learning

#### 4.6.1. Validation of data and method

We used a zero-shot decoding approach (Palatucci et al., 2009) to validate the data and method with respect to previously reported results. Regularized multivariate ridge regression, as implemented in scikit-learn (Pedregosa et al., 2011), was used to fit a model that predicts the semantic feature vector of a target concept based on brain response. The sensor-level MEG responses were first standardized with respect to concepts, such that the mean of each predictor (time point-channel pair) was 0 and the standard deviation was 1. We used leave-one-out cross validation, such that models were trained on 59 out of the 60 concepts and evaluated on the remaining concept the model had not been trained on, for all permutations. To evaluate performance, we calculated the cosine similarity between the prediction and the target vector and calculated its rank in relation to 99 other Finnish nouns with high frequency, high concreteness (as recorded by the ‘physical entity’ property in Finnish WordNet (Lindén & Niemi, 2014)), and similar word length to the present stimuli. Details of the validation set are in Appendix B. We then calculated the mean rank accuracy (Markowitz et al., 2017) as (1 *−* mean(*R*_*t*_)*/N*) *×* 100%, where *R*_*t*_ is the rank of the target vector (in relation to the 99 other vectors) with respect to cosine similarity, and *N* = 100, the number of vectors compared. Statistical significance was evaluated using a permutation test with 1000 iterations as in Kivisaari et al. (2019). At each iteration brain data of a randomly selected participant was shuffled and decoding was performed. *p*-values were calculated by the proportion of simulated mean rank accuracy values that were at least as high as the true mean rank accuracy.

#### 4.6.2. Mapping brain response to semantic space as a function of time

In accordance with Carlson et al. (2013) and Grootswagers et al. (2017), we performed temporal cross-decoding, in which models are trained and tested on different time windows to check whether there is information generalization in the brain across different time points. For this we trained and tested models on pairs of 20 ms time windows on data averaged over all participants. To explore the progression of semantic understanding in more detail we compared two types of models on average sensor-level MEG responses. First, we used sliding windows of fixed length similar to Sudre et al. (2012) and Rupp et al. (2017). Second, we developed a method to examine cumulatively widening windows (see details below). In both cases, we evaluated models based on prediction-target cosine similarity (higher similarity indicates that the model better predicts the target word). In comparison to performance based on rank, performance based on similarity is more sensitive and allows to capture the dynamics of semantic understanding even for single concepts in a finer-grained manner. Both types of models were evaluated using leave-one-out cross validation. Significance testing to compare the models was performed using a permutation test which compared the two models at each time point. Again, 1000 iterations with shuffled data were performed and the *p*-values for each time point were calculated from the proportion of differences that were at least as large as the observed difference.

##### Sliding fixed-length window

Regression models were trained and tested on fixed-length subsets of MEG data. A sliding window of 20 ms was used, with no overlap between adjacent windows. Each subset was evaluated independently of others. Similar models have been used by Hultén et al. (2021); Rupp et al. (2017); Sudre et al. (2012) and we expected a pattern of gradual increase, followed by a decrease, in cosine similarity.

##### Cumulative window

Regression models were trained and tested on cumulative subsets of MEG data. The window size was sequentially increased by 20 ms. Thus, all previously encoded information was included in the estimation and model evaluation (Figure 1B).

#### 4.6.3. Identifying individual differences

To compare the progression of semantic understanding between participants, we used the cumulative models. We first calculated the progression of semantic information for all concepts for each participant. We then focused on the point of plateau, the time point after which there was less than 5% increase in prediction-target similarity. Concepts that had their plateaus already at the first time point or did not reach plateau by the last point were excluded (mean: 4, range: 2-11 concepts out of 60 removed per participant). We compared these plateau points using a linear mixed model, with random intercepts for concepts.

### 4.7. Representational Similarity Analysis

We performed Representational Similarity Analysis (RSA; Kriegeskorte, 2008) on source localized MEG data to extract the brain areas accounting for highest similarity between the brain activation patterns and vector representations of the words for different time windows. This was done using the MNE-RSA software package (van Vliet, 2022). RSA was performed for both sliding and cumulative time windows.

The model dissimilarity matrix (DSM) was obtained by calculating pairwise cosine distances between vector representations of the words. This was followed by the calculation of brain DSMs for each participant at each source-level vertex with a searchlight patch radius of 2 cm for the time window of interest. Brain DSMs were computed by calculating pairwise Pearson correlation coefficient between brain signals in response to different stimuli. The relationship between the brain DSMs and model DSMs was quantified by Spearman rank correlation coefficients for each participant. This resulted in participant-level RSA maps (Figure 4A and B, top rows). We then used a cluster permutation test (Maris & Oostenveld, 2007) across participants with a cluster threshold of *p* = .01, a cluster-wide significance threshold of *p* = .05 and 5000 permutations in accordance with Hultén et al. (2021).

## 5. Acknowledgments

We would like to thank Jenna Kanerva and Filip Ginter at the University of Turku for development of the Finnish language word2vec model. We also thank Ali Faisal for assistance in the early stages of the project and Anni Nora for helpful comments on a draft. Computational resources were provided by the Aalto Science-IT project. This research was funded by the Academy of Finland (#315553 to RS, #310988 and #343385 to MvV, #286070 to SK, #287474 to AH) and the Sigrid Jusélius Foundation (to RS).

---

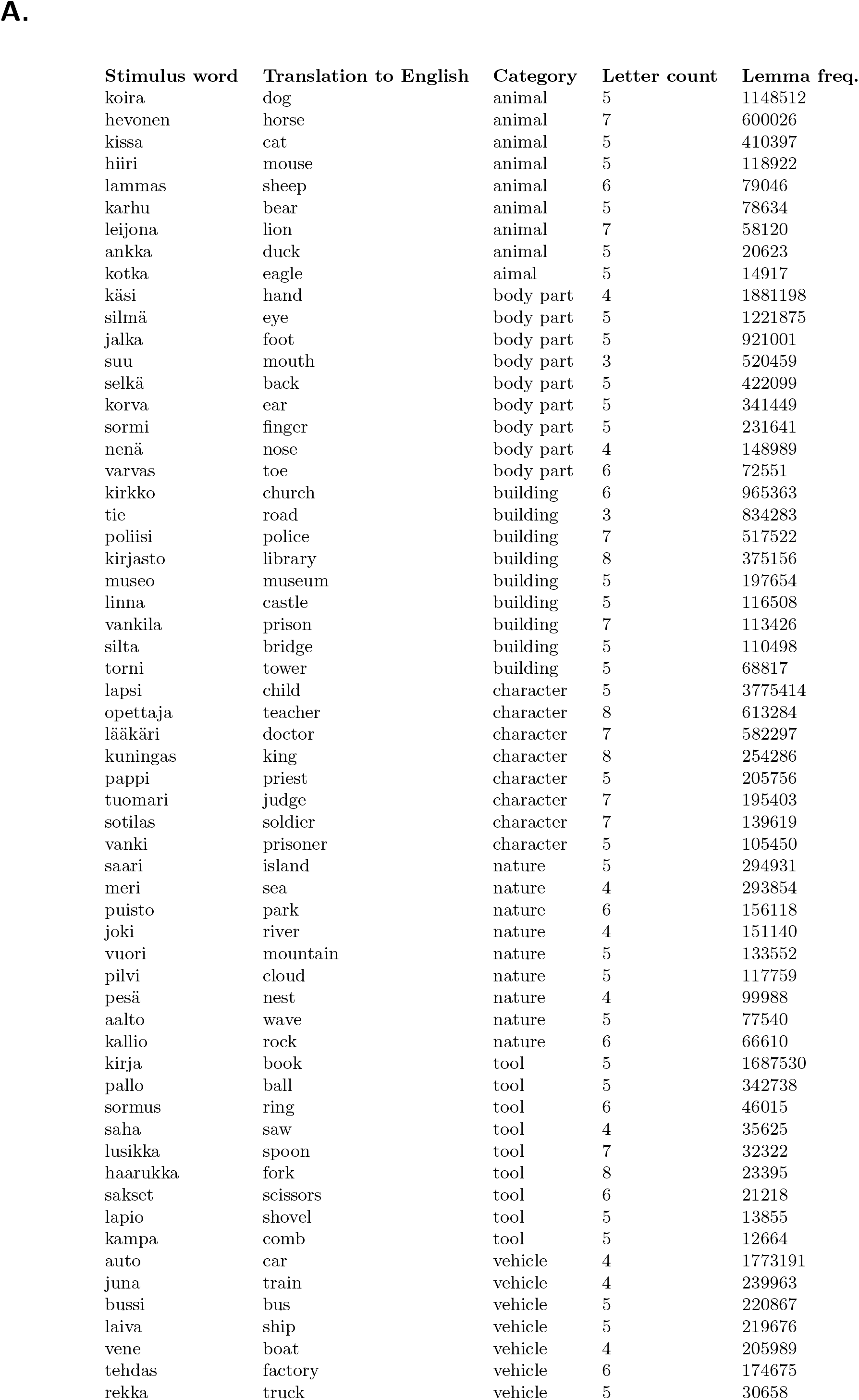
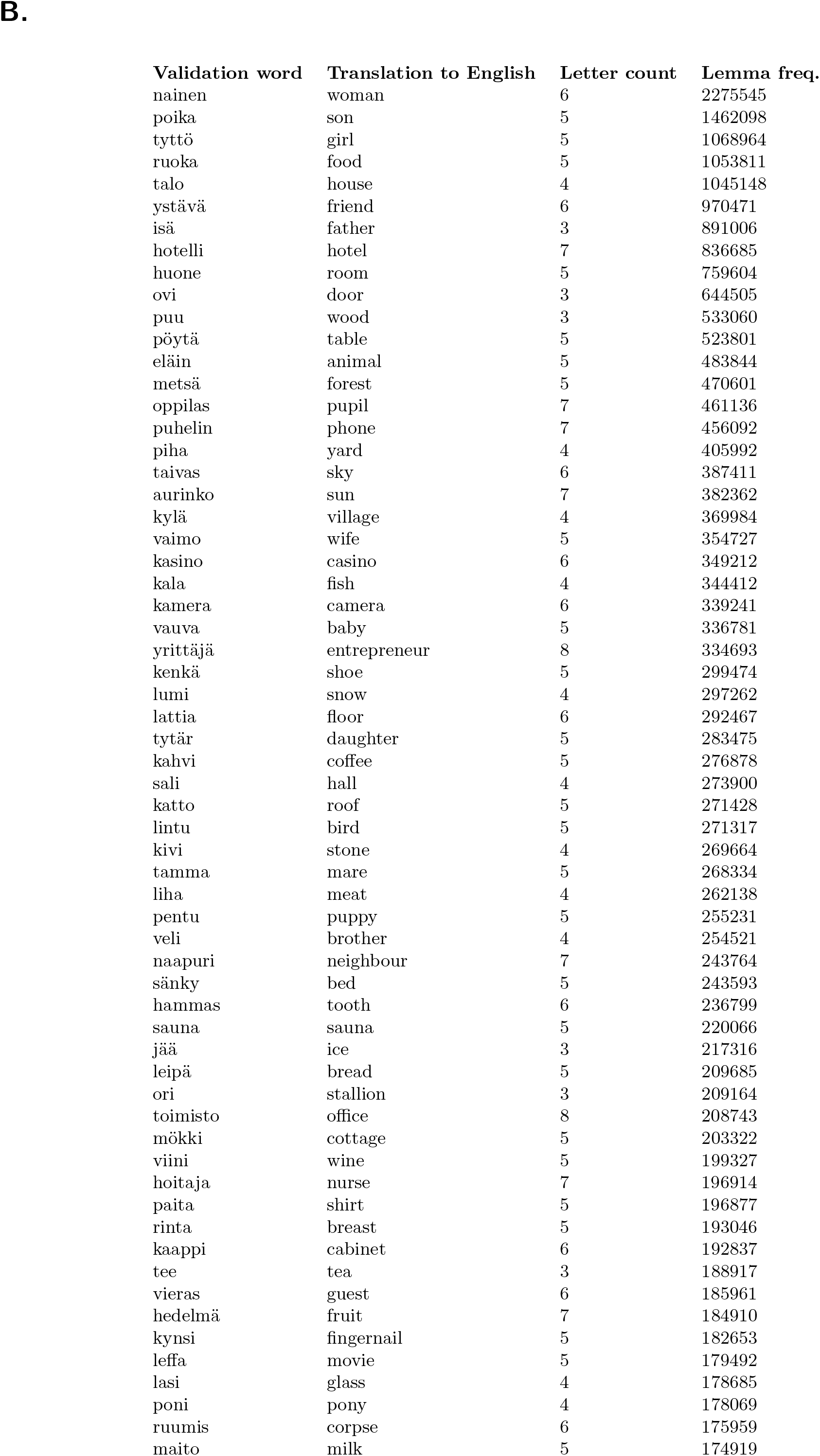
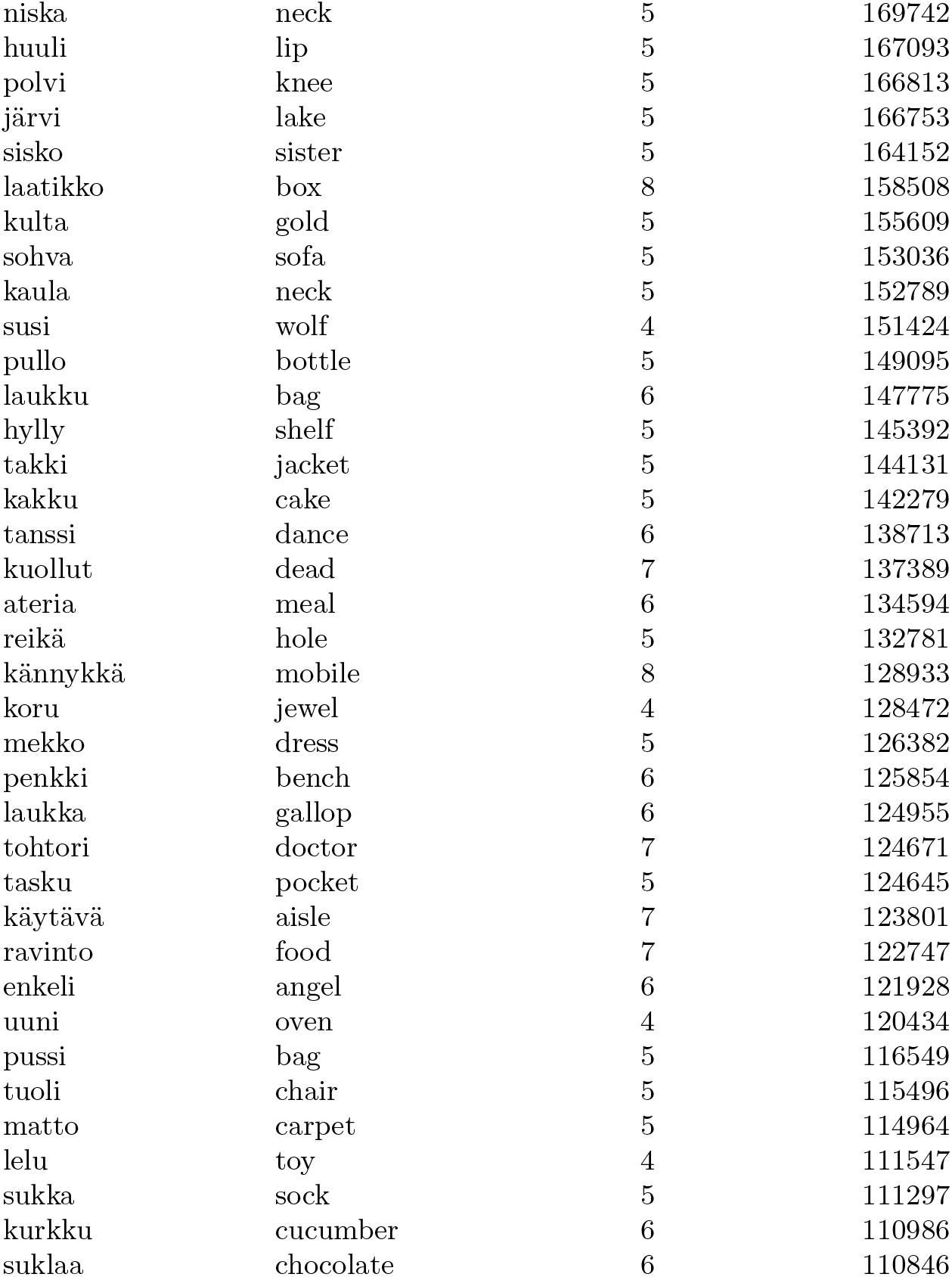

| Stimulus word | Translation to English | Category | Letter count | Lemma freq. |
| --- | --- | --- | --- | --- |
| koira | dog | animal | 5 | 1148512 |
| hevonen | horse | animal | 7 | 600026 |
| kissa | cat | animal | 5 | 410397 |
| hiiri | mouse | animal | 5 | 118922 |
| lammas | sheep | animal | 6 | 79046 |
| karhu | bear | animal | 5 | 78634 |
| leijona | lion | animal | 7 | 58120 |
| ankka | duck | animal | 5 | 20623 |
| kotka | eagle | animal | 5 | 14917 |
| käsi | hand | body part | 4 | 1881198 |
| silmä | eye | body part | 5 | 1221875 |
| jalka | foot | body part | 5 | 921001 |
| suu | mouth | body part | 3 | 520459 |
| selkä | back | body part | 5 | 422099 |
| korva | ear | body part | 5 | 341449 |
| sormi | finger | body part | 5 | 231641 |
| nenä | nose | body part | 4 | 148989 |
| varvas | toe | body part | 6 | 72551 |
| kirkko | church | building | 6 | 965363 |
| tie | road | building | 3 | 834283 |
| poliisi | police | building | 7 | 517522 |
| kirjasto | library | building | 8 | 375156 |
| museo | museum | building | 5 | 197654 |
| linna | castle | building | 5 | 116508 |
| vankila | prison | building | 7 | 113426 |
| silta | bridge | building | 5 | 110498 |
| torni | tower | building | 5 | 68817 |
| lapsi | child | character | 5 | 3775414 |
| opettaja | teacher | character | 8 | 613284 |
| lääkäri | doctor | character | 7 | 582297 |
| kuningas | king | character | 8 | 254286 |
| pappi | priest | character | 5 | 205756 |
| tuomari | judge | character | 7 | 195403 |
| sotilas | soldier | character | 7 | 139619 |
| vanki | prisoner | character | 5 | 105450 |
| saari | island | nature | 5 | 294931 |
| meri | sea | nature | 4 | 293854 |
| puisto | park | nature | 6 | 156118 |
| joki | river | nature | 4 | 151140 |
| vuori | mountain | nature | 5 | 133552 |
| pilvi | cloud | nature | 5 | 117759 |
| pesä | nest | nature | 4 | 99988 |
| aalto | wave | nature | 5 | 77540 |
| kallio | rock | nature | 6 | 66610 |
| kirja | book | tool | 5 | 1687530 |
| pallo | ball | tool | 5 | 342738 |
| sormus | ring | tool | 6 | 46015 |
| saha | saw | tool | 4 | 35625 |
| lusikka | spoon | tool | 7 | 32322 |
| haarukka | fork | tool | 8 | 23395 |
| sakset | scissors | tool | 6 | 21218 |
| lapio | shovel | tool | 5 | 13855 |
| kampa | comb | tool | 5 | 12664 |
| auto | car | vehicle | 4 | 1773191 |
| juna | train | vehicle | 4 | 239963 |
| bussi | bus | vehicle | 5 | 220867 |
| laiva | ship | vehicle | 5 | 219676 |
| vene | boat | vehicle | 4 | 205989 |
| tehdas | factory | vehicle | 6 | 174675 |
| rekka | truck | vehicle | 5 | 30658 |

B.
| Validation word | Translation to English | Letter count | Lemma freq. |
| --- | --- | --- | --- |
| nainen | woman | 6 | 2275545 |
| poika | son | 5 | 1462098 |
| tyttö | girl | 5 | 1068964 |
| ruoka | food | 5 | 1053811 |
| talo | house | 4 | 1045148 |
| ystävä | friend | 6 | 970471 |
| isä | father | 3 | 891006 |
| hotelli | hotel | 7 | 836685 |
| huone | room | 5 | 759604 |
| ovi | door | 3 | 644505 |
| puu | wood | 3 | 533060 |
| pöytä | table | 5 | 523801 |
| eläin | animal | 5 | 483844 |
| metsä | forest | 5 | 470601 |
| oppilas | pupil | 7 | 461136 |
| puhelin | phone | 7 | 456092 |
| piha | yard | 4 | 405992 |
| taivas | sky | 6 | 387411 |
| aurinko | sun | 7 | 382362 |
| kylä | village | 4 | 369984 |
| vaimo | wife | 5 | 354727 |
| kasino | casino | 6 | 349212 |
| kala | fish | 4 | 344412 |
| kamera | camera | 6 | 339241 |
| vauva | baby | 5 | 336781 |
| yrittäjä | entrepreneur | 8 | 334693 |
| kenkä | shoe | 5 | 299474 |
| lumi | snow | 4 | 297262 |
| lattia | floor | 6 | 292467 |
| tytär | daughter | 5 | 283475 |
| kahvi | coffee | 5 | 276878 |
| sali | hall | 4 | 273900 |
| katto | roof | 5 | 271428 |
| lintu | bird | 5 | 271317 |
| kivi | stone | 4 | 269664 |
| tamma | mare | 5 | 268334 |
| liha | meat | 4 | 262138 |
| pentu | puppy | 5 | 255231 |
| veli | brother | 4 | 254521 |
| naapuri | neighbour | 7 | 243764 |
| sänky | bed | 5 | 243593 |
| hammas | tooth | 6 | 236799 |
| sauna | sauna | 5 | 220066 |
| jää | ice | 3 | 217316 |
| leipä | bread | 5 | 209685 |
| ori | stallion | 3 | 209164 |
| toimisto | office | 8 | 208743 |
| mökki | cottage | 5 | 203322 |
| viini | wine | 5 | 199327 |
| hoitaja | nurse | 7 | 196914 |
| paita | shirt | 5 | 196877 |
| rinta | breast | 5 | 193046 |
| kaappi | cabinet | 6 | 192837 |
| tee | tea | 3 | 188917 |
| vieras | guest | 6 | 185961 |
| hedelmä | fruit | 7 | 184910 |
| kynsi | finger nail | 5 | 182653 |
| leffa | movie | 5 | 179492 |
| lasi | glass | 4 | 178685 |
| poni | pony | 4 | 178069 |
| ruumis | corpse | 6 | 175959 |
| maito | milk | 5 | 174919 |
| niska | neck | 5 | 169742 |
| huuli | lip | 5 | 167093 |
| polvi | knee | 5 | 166813 |
| järvi | lake | 5 | 166753 |
| sisko | sister | 5 | 164152 |
| laatikko | box | 8 | 158508 |
| kulta | gold | 5 | 155609 |
| sohva | sofa | 5 | 153036 |
| kaula | neck | 5 | 152789 |
| susi | wolf | 4 | 151424 |
| pullo | bottle | 5 | 149095 |
| laukku | bag | 6 | 147775 |
| hylly | shelf | 5 | 145392 |
| takki | jacket | 5 | 144131 |
| kakku | cake | 5 | 142279 |
| tanssi | dance | 6 | 138713 |
| kuollut | dead | 7 | 137389 |
| ateria | meal | 6 | 134594 |
| reikä | hole | 5 | 132781 |
| kännykkä | mobile | 8 | 128933 |
| koru | jewel | 4 | 128472 |
| mekko | dress | 5 | 126382 |
| penkki | bench | 6 | 125854 |
| laukka | gallop | 6 | 124955 |
| tohtori | doctor | 7 | 124671 |
| tasku | pocket | 5 | 124645 |
| käytävä | aisle | 7 | 123801 |
| ravinto | food | 7 | 122747 |
| enkeli | angel | 6 | 121928 |
| uuni | oven | 4 | 120434 |
| pussi | bag | 5 | 116549 |
| tuoli | chair | 5 | 115496 |
| matto | carpet | 5 | 114964 |
| lelu | toy | 4 | 111547 |
| sukka | sock | 5 | 111297 |
| kurkku | cucumber | 6 | 110986 |
| suklaa | chocolate | 6 | 110846 |

